# Next-generation CRISPR gene-drive systems using Cas12a nuclease

**DOI:** 10.1101/2023.02.20.529271

**Authors:** Sara Sanz Juste, Emily M. Okamoto, Xuechun Feng, Victor López Del Amo

## Abstract

One method for reducing the impact of vector-borne diseases is through the use of CRISPR-based gene drives, which manipulate insect populations due to their ability to rapidly propagate desired genetic traits into a target population. However, all current gene drives employ a Cas9 nuclease that is constitutively active, impeding our control over their propagation abilities and limiting the generation of novel gene drive arrangements. Yet, other nucleases such as the temperature-sensitive Cas12a have not been explored for gene drive designs. To address this, we herein present a proof-of-concept gene-drive system driven by Cas12a that can be regulated via temperature modulation. Furthermore, we combined Cas9 and Cas12a to build double gene drives capable of simultaneously spreading two independent engineered alleles. The development of Cas12a-mediated gene drives provides an innovative option for designing next-generation vector control strategies to combat disease vectors and agricultural pests.

## INTRODUCTION

One strategy for controlling vector-borne pathogens and agricultural pests involves the use of CRISPR gene-drive systems, which represent a novel genetic tool for propagating desired traits into wild insect populations^1^. The engineered genetic elements that comprise gene drives are rapidly disseminated through a population because they are self-propagating and can bias their inheritance rate. Current CRISPR-based gene drives consist of a three-component transgene: (i) Cas9, a nuclease that produces DNA double-strand breaks; (ii) a guide RNA (gRNA) that leads Cas9 to cleave the DNA at a target site; and (iii) two homology arms matching both sides of the cut site to promote homology-directed repair (HDR). Gene drives propagate by first generating a DNA double strand break in the wildtype allele, which is repaired by HDR using the gene-drive containing chromosome as a template, ultimately resulting in the original wildtype allele being replaced by that of a gene-drive one^2^. At the population level, when a gene-drive engineered individual encounters a wildtype one, the subsequent allelic conversion promotes biased Mendelian inheritance (50%) towards super-Mendelian inheritance (>50%), allowing rapid gene drive propagation through the population. In fact, mosquitoes with CRISPR gene-drive modifications have been observed to reach nearly 100% inheritance of the desired allele in consecutive generations^3–6^.

However, even though these drives can propagate desirable traits for controlling disease spread into a native population, gene-drive propagation is difficult to control due to the always-active nature of the Cas9 nuclease. To develop some layers of control over the self-propagation of gene drives, multiple strategies have been developed^1^. For example, a split gene drive was built by placing the Cas9 and gRNA components at two different loci into two different transgenic lines; here, the gene drive was activated only when these two strains were combined by genetic crosses^7,8^. Yet, the generation of multiple transgenic lines, especially for organisms with difficulties in obtaining high transgenic efficiencies such as mosquitoes^9,10^ could be an issue. To fine-tune gene drives using a small molecule, a drug-regulated system was developed, but this approach required a less efficient modified Cas9 and a more complex genetic circuitry^11,12^. Lastly, ECHACR, ERACR, and anti-CRISPR approaches could also help control gene-drive activities^13,14^, but these systems may not be preferable as they require a secondary release of genetically-modified animals, imposing potential regulatory issues.

Furthermore, all current gene-drive methods employ a Cas9 nuclease^1^, limiting the creation of novel arrangements towards next-generation gene-drive systems. Yet, other nucleases, such as Cas12a, have demonstrated efficient genome editing in a broad range of organisms^15–18^. In *Drosophila melanogaster*, Cas12a-based genome editing produced efficient gene disruption^19,20^; however, the DNA homology-directed repair (HDR) triggered by Cas12a has yet not been explored in insects.

Based on previous works regarding Cas12a-involved genome editing, we hypothesized that the Cas12a-mediated gene drive should be a suitable strategy for addressing the lack of control over Cas9 activity while increasing the available toolkit for future genetic strategies for vector control. First, a Cas12a gene-drive system could be modulated with temperature, which would increase our control over its self-propagation, thereby adding an extra level of containment. In principle, Cas12a gene drives should remain inactive at low temperatures while activating their self-propagation abilities at higher temperatures. Second, a functional Cas12a gene-drive system would be orthogonal to a Cas9 one, which should open new avenues toward developing next-generation arrangements such as double-drive systems, where two gene-drive elements (driven by distinct nucleases) spread concurrently. For these reasons, we wanted to explore the use of Cas12a in a gene-drive setting.

Thus, we herein explored the feasibility of a temperature-regulated Cas12a gene-drive system in *Drosophila melanogaster*. We first present a Cas12a gene-drive system that promotes super-Mendelian inheritance in *Drosophila melanogaster*, revealing that Cas12a can trigger efficient HDR in the germline in a temperature-dependent manner. In addition, we also built a double gene drive system by placing Cas9 and Cas12 together and demonstrated that two independent genetic elements could spread concurrently within the same organism. This work presents the first Cas12a-based gene-drive system in insects while bringing new opportunities for developing novel gene drives to control disease vectors and agricultural pests.

## RESULTS

### A Cas12a gene-drive system induces super-Mendelian inheritance in a temperature-dependent manner

To determine if the Cas12a nuclease could generate a functional gene drive, we employed a CopyCat gRNA-only gene-drive strategy^11,21^ using *Drosophila melanogaster*. Briefly, a CopyCat gene-drive element carries a gRNA gene and two homology arms neighboring the gRNA cut site, allowing the gRNA cassette to copy itself onto the homologous chromosome. The CopyCat transgene propagates only when combined with a Cas transgene, which itself is inherited in a Mendelian fashion (50%) (**Fig. 1a**)^11,21^.

To test the Cas12a-based gene drives, we built three different transgenic lines: i) one carrying a Cas12a cassette inserted into the yellow(y) locus (X chromosome [Chr.X]) and marked with DsRed; ii) a CopyCat transgenic line carrying the *e1*-gRNA under the control of the *Drosophila* U6:3 promoter and Opie2-GFP inserted in the *ebony* locus (Chr.III) for tracking the transgene; and iii) another CopyCat transgenic line carrying the *e4*-gRNA under the control of *Drosophila* U6:3 promoter and Opie2-GFP inserted in the *ebony* locus (Chr.III) (**Supplementary Fig. 1a**). It is important to note that our Cas12a-based CopyCat gene drives were built such that both homology arms cover the five–nucleotide-overhang staggered cuts from the Cas12a nuclease, so the homology arms have five overlapping nucleotides on their respective ends (**Supplementary Fig.1b**).

To show the feasibility of the Cas12a-mediated gene drive, we first evaluated our system without temperature control by keeping the flies at the standard incubation temperature of 25°C. Next, we crossed Cas12a transgenic males to the CopyCat gene-drive transgenic females (either *e1*-gRNA or *e4*-gRNA) (**Fig. 1b**; F0 generation). Then, we collected F1 transheterozygous females carrying both Cas12a and CopyCat transgenes (**Fig. 1b**; F1) and performed single-pair crossings to *ebony* mutants (e-/e-) to evaluate super-Mendelian inheritance via the GFP marker of the gene-drive element in the F2 progeny (**Fig. 1b**). If the gene drive is active, the GFP marker should score inheritance rates higher than 50%. Indeed, we observed super-Mendelian inheritance in both cases, with the *e1*-gene drive (*e1*-GD) and the *e4*-GD element showing inheritance rates of 83% and 67%, respectively (**Fig. 1c; Supplementary Data 1**). We also scored the inheritance of the Cas12a (marked with DsRed) in both cases and confirmed its expected Mendelian inheritance of 50% (**Supplementary Data 1**).

To further investigate whether temperature could regulate the super-Mendelian rates displayed by our gene-drive elements, we followed the same approach as above (**Fig. 1b)** and repeated the experiments at 18°C and 29°C. As we would hope for a system with reduced activity at lower temperatures, we observed reduced super-Mendelian inheritance rates of 73% (*e1*-GD) and 52% (*e4*-GD) at 18°C (**Fig. 1c**), and increased inheritance rates of 84% (*e1*-GD) and 81% (e4-GD) at 29°C. Indeed, these results indicate the temperature-dependent behavior of our Cas12a gene-drive system, providing new opportunities to control gene drive activity (**Fig. 1c; Supplementary Data 1**).

**Fig.1.**
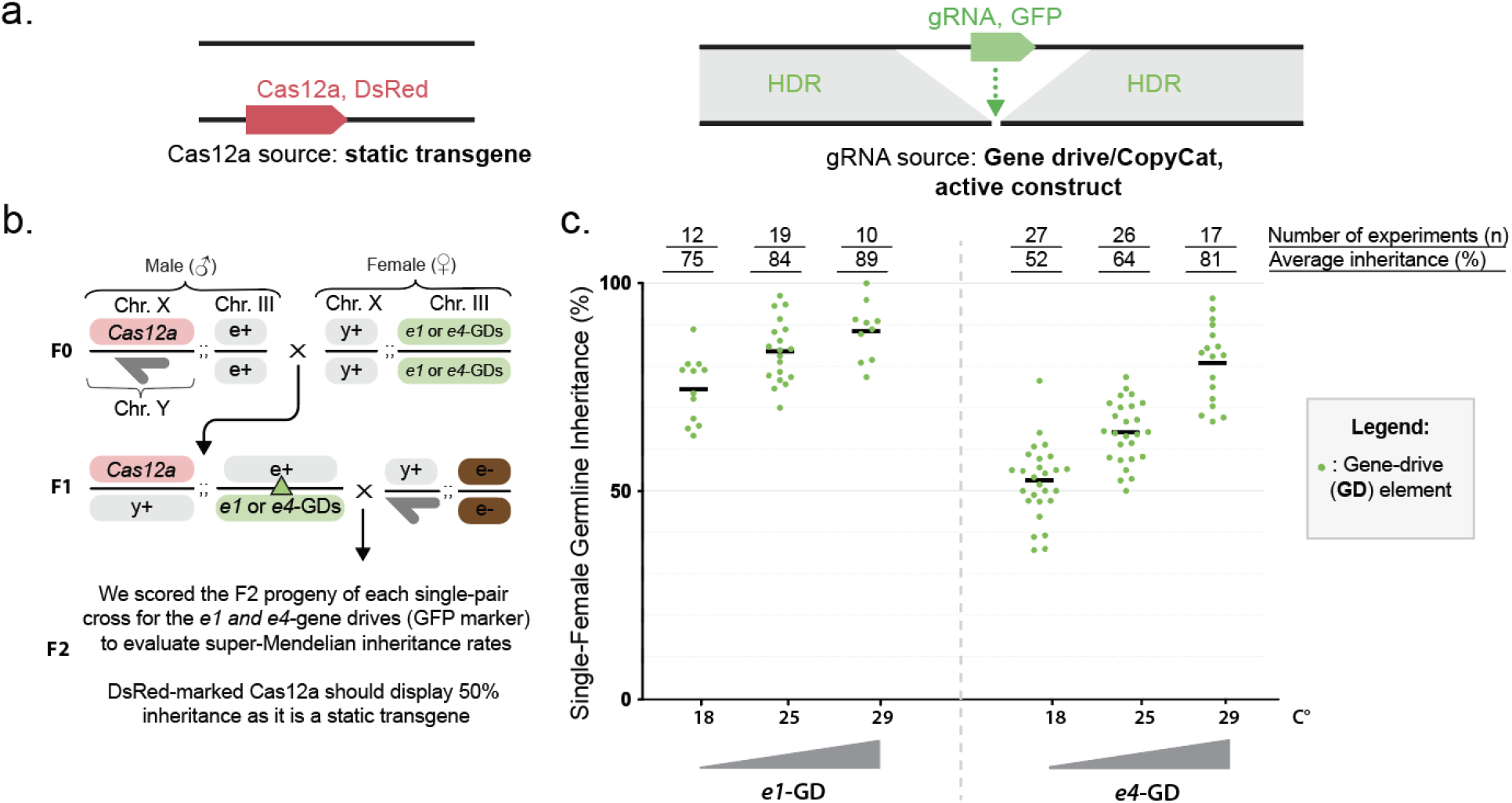
Cas12a-based gene drives display super-Mendelian inheritance rates modulated by temperature. **(a)** Schematic of the CopyCat gene-drive system. The DsRed-marked Cas12a is a static transgene that provides the nuclease for copying the GFP-marked CopyCat element by allelic conversion, which is driven by the surrounding homology arms. **(b)** Cross scheme of males expressing Cas12a crossed to virgin females carrying the ebony CopyCat constructs (*e1* or *e4* gene drives). Collected virgin females (Cas12a-dsRed + gene drive-GFP) were crossed to ebony mutant males to score germline transmission rates by screening the GFP marker in the F2 progeny. The dark-gray half arrow indicates the male Y chromosome. The green triangle in the F1 female indicates potential gene drive copying onto the wildtype chromosome. **(c)** Assessment of gene-drive activity in the germline of F1 females by phenotypically scoring the F2 progeny for the GFP-marked ebony CopyCat constructs. Measurements of inheritance rates are reported on top of the graph along with the average inheritance (%) (also as black bars on the graph) and the number of F1 crosses performed (n).

As the *ebony* gene is a recessive locus, our experimental design made it possible to obtain estimated conversion rates/HDR, formation rates for indels/resistant alleles, and wildtype/uncut rates for our gene-drive elements based on phenotypes and GFP presence (**Supplementary Fig.2**). At the standard temperature of 25°C, we observed a conversion rate of 91% and 93% for *e1*- and *e4*-GD, respectively, indicating overall high HDR efficiency of the Cas12a-mediated gene drive. In line with this high conversion rate, we only observed the formation of resistant alleles in 8% (*e1*-GD) and 5% (*e4*-GD) of gene-drive individuals. Lastly, we detected 9% and 29% uncut alleles for our *e1*-GD and *e4*-GD at 25°C, respectively (**Supplementary Fig.2; Supplementary Data 1**). This indicates that the gene-drive elements do not have 100% cutting rates and suggests that the *e1*-GD is more active than the *e4*-GD at 25°C, explaining the observed inheritance differences between the gene-drive elements. Overall, we observed an increase in super-Mendelian inheritance rates when experiments were performed at higher temperatures with both alleles (*e1*-GD and *e4*-GD). Additionally, *e4*-GD showed no gene-drive activity at 18°C, providing a new temperature-sensitive option for controlling these active genetic elements (**Fig.1c; Supplementary Data 1**).

### Cas9 and Cas12a promote the propagation of two independent gene drives within the same organism

To show the feasibility of two independent gene drives spreading simultaneously in a single organism, we employed a double CopyCat gene-drive strategy. The two different Cas nucleases (Cas9 and Cas12a) were inserted individually in different chromosomes as static transgenes: Cas9 was inserted in Chr. III and Cas12a was inserted in Chr. X. (**Fig. 2a**). For the CopyCat gene-drive elements, we combined the *w2*-gRNA (*w2*-GD) with Cas9 to target the *white* gene in Chr. X.; this gRNA activity was already validated in our previous study^7,11^.

**Fig. 2.**
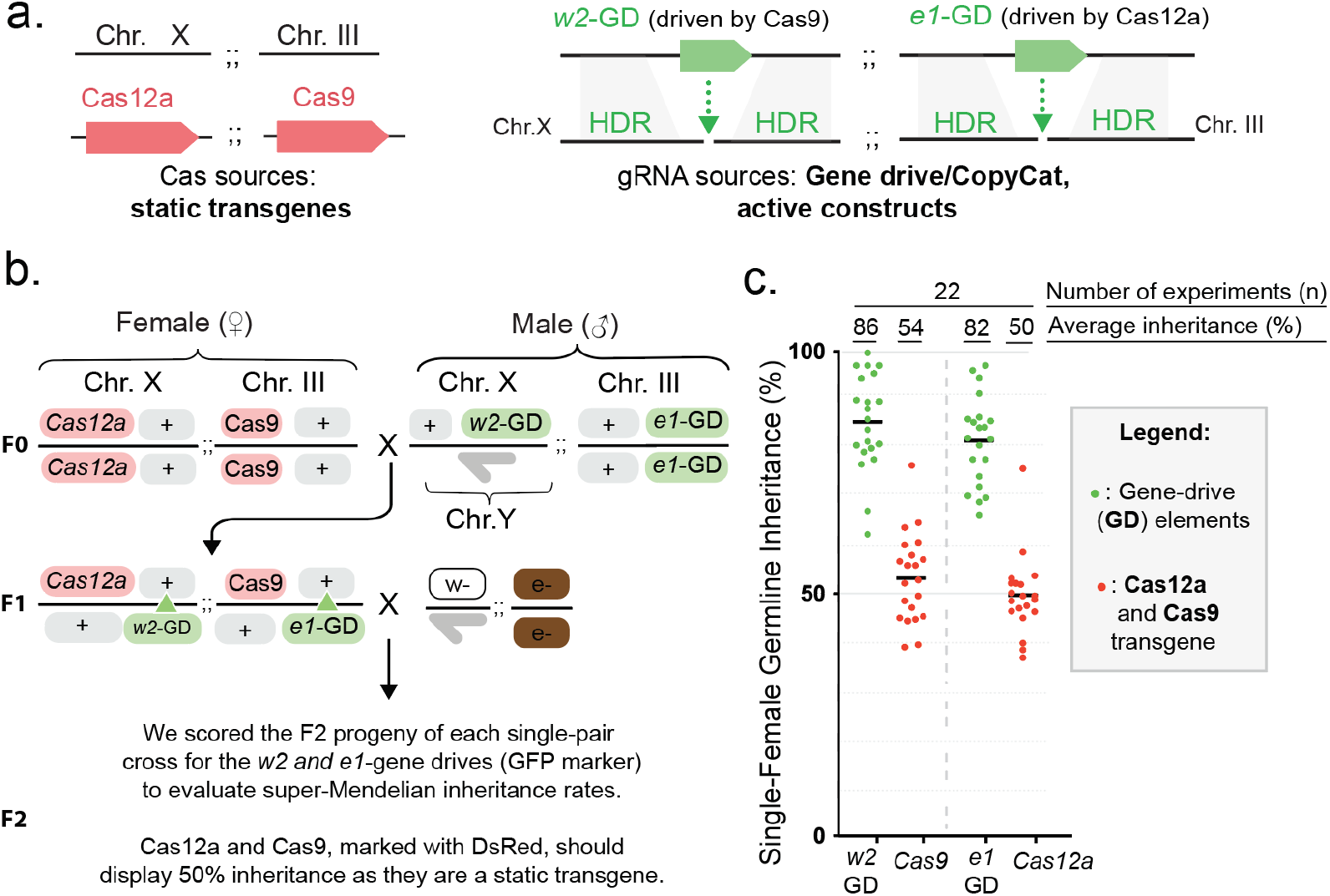
Double gene drives driven by Cas9 and Cas12a display simultaneous super-Mendelian inheritance rates of two independent genetic elements. **(A)** Schematic of the double CopyCat gene-drive system. The DsRed-marked Cas12a and Cas9 represent the nuclease sources as static transgenes. Two independent CopyCat gene-drive elements (*w2*-GD and *e1*-GD) are driven by the Cas9 or Cas12a nucleases. To track their inheritance, we marked all four transgenes without overlapping fluorescent markers in any body parts. **(B)** Cross scheme of males expressing Cas12a and Cas9 crossed to virgin females carrying two CopyCat constructs (*e1* and *w2* gene drives). Collected F1 virgin females (Cas12a-Cas9-DsRed + gene drives-GFP) were crossed to ebony and white mutant males to score germline transmission rates by screening the GFP markers in the F2 progeny. A dark-gray half arrow indicates the male Y chromosome. The green triangles in the F1 female indicate potential gene drive copying onto the wildtype allele. **(C)** Evaluation of gene-drive activity in the germline of F1 females by phenotypically scoring the F2 progeny of the crosses for the GFP-marked gene-drive constructs. Our measurements of inheritance rates are reported on top of the graph along with the inheritance average (%) (also as black bars on the graph) and the number of F1 crosses performed (n).

For the second gene-drive element, we combined the *e1*-GD with the Cas12a nuclease to target the ebony gene (**Fig. 2a**); the *e1*-gRNA was validated in our results in **Figure 1**. To allow independent tracking when scoring, we marked all transgenes with various fluorescent markers located in different fly body parts. Cas12a and Cas9 were marked with DsRed in the eye and thorax, respectively, and the gRNA-only gene drives driven by Cas12a (*e1*-GD) and Cas9 (*w2*-GD) were marked with GFP in the abdomen and eye, respectively (**Supplementary Fig.1a**).

To assess their inheritance bias, we crossed Cas12a/Cas9 females with the gene-drive transgenic males containing the two above mentioned gene-drive elements. Subsequently, we collected F1 females containing all four transgenes and crossed them to white and ebony mutant males to score transgene inheritance in the F2 progeny (**Fig. 2b**). As expected, we observed Mendelian inheritance rates for the Cas9 and Cas12a nucleases (**Fig. 2c**; **Supplementary Data 2**) and an 86% average inheritance rate for w2-GD (driven by Cas9) and an 81% average inheritance rate for *e*1-GD (driven by Cas12a) in F2 offspring (**Fig. 2c**). These results indicate that two independent gene-drive cassettes driven by distinct nucleases can display simultaneous super-Mendelian inheritance of genetically engineered elements in the same organism.

## DISCUSSION

This manuscript presents a proof-of-concept for a temperature-controlled Cas12a gene drive and demonstrates the feasibility of two gene drives driven by two different nucleases (Cas9 and Cas12a) propagating in a single organism. We showed that Cas12a could induce super-Mendelian inheritance in two different genetically engineered alleles (*e1*-GD and *e4*-GD) in a temperature-dependent manner. Both gene drives, driven by validated gRNAs targeting the *ebony* gene for disruption^20^, displayed the desirably lower super-Mendelian rates at 18°C and increased gene-drive efficiency at 25°C and 29°C. Our results demonstrate that Cas12a-based HDR, which has been extensively used in other organisms^17,18,22^, can also be achieved in a gene drive context using *Drosophila melanogaster*. This work shows the functionality of a Cas12a gene-drive system triggering super-Mendelian inheritance rates in an insect germline, and the first presenting a dual gene-drive in any organism, both advances addressing concerns in Cas9-based vector control strategies.

Interestingly and in line with the activity of Cas12a at lower temperatures, we did not observe super-Mendelian inheritance rates when the *e4*-GD experiments were performed at 18°C, indicating that Cas12a-based gene-drive inactivation may be possible by switching to lower temperatures. However, it is important to mention that the *e1*-GD still displayed super-Mendelian inheritance rates at 18°C. This observation could be explained by differences in gRNA activities as distinct gRNAs promoted variable levels of disruption when targeting the *ebony* gene^20^.

Altogether, our experiments show that the use of a Cas12a nuclease can add a temperature-dependent layer of containment over gene-drive propagation. In addition to providing some level of inactivity when desired, unlike to the always-on Cas9 nuclease, gene-drive containment is an important feature demanded by the gene drive community and regulators^23–26^. Also, these Cas12a-based gene-drive systems displayed super-Mendelian inheritance rates comparable to previous Cas9-based systems (80–90%) that efficiently spread in a target population under laboratory conditions^27^, though future investigations are needed to explore the propagation dynamics of Cas12a-based systems and the formation of resistant alleles that are also a concern for gene-drive propagation^28,29^. Interestingly, Cas12a-introduced DNA breaks are far away from the DNA recognition site (PAM) compared to Cas9. In fact, mutations containing small indels far away from the PAM site were observed in plants and zebrafish^17,30^, and this may be harnessed to improve gene drive propagation, as resistance alleles far away from the PAM site could still be recognized by the Cas12a-based gene drive to allow its full introgression into a target population.

In addition, a Cas12a system should improve current Cas9-based genetic sterile insect technique (gSIT) that involves high maintenance costs from rearing two separated stocks (Cas9 and gRNAs) and sexing of the separated strains to generate sterile males through genetic crosses^31,32^. By using a Cas12a nuclease instead, a single strain containing Cas12a and gRNAs together could be generated and kept as a regular stock at 18°C, since Cas12a should be inactive. Then, sterile males would be generated by switching the stock to higher temperatures^33^ while avoiding the crossing step required in Cas9 settings. This could potentially reduce logistical issues when high throughput rearing is needed (i.e., for field release purposes).

Once we demonstrated the feasibility of a gene-drive approach using Cas12a, we also questioned whether the propagation of two independent gene-drive elements driven by two different nucleases is possible in the same organism. The combination of two transgenic lines, one containing Cas9 and Cas12a and a second line containing two gRNA-only drive elements driven by each of the nucleases, showed super-Mendelian inheritance rates for *w2*-GD (86%) and *e1*-GD (82%) at 25°C. Both gene-drive elements showed comparable though slightly lower inheritance rates when tested singularly. *e1*-GD had comparable inheritance rates: 84% singularly and 82% when combined with the *w2*-GD. We previously tested the singular *w2*-GD driven by Cas9 and saw 90% super-Mendelian inheritance rates^11^, which is slightly higher than the 86% inheritance rate observed in our double gene-drive system. These differences in inheritance rates are most likely due to the different genomic locations of Cas9 and/or the promoter employed in each situation, as we previously reported. In this work, we utilized a Cas9 located in the third chromosome driven by the *nanos* promoter, whereas when *w2*-GD was tested singularly before, it was driven by a Cas9 inserted in the X chromosome driven by the *vasa* promoter^7^.

Regardless, this double gene-drive system promoting super-Mendelian inheritance rates of two independent transgenes opens a new avenue for designing innovative approaches with two distinct nucleases. For example, this could be implemented to design new insect-vector population suppression systems such as the proposed double drives, which target two different genomic locations^34^. A double drive system, in which Cas9 and Cas12a would target sex-determination genes at two locations (within the same gene or at distinct loci), could improve gene drive spread, as indels generated at one target site would not affect the propagation of the other gene drive cassette, and ultimately ensure its full propagation while improving current suppression strategies. It is important to note that Cas9 and Cas12a have different PAM requirements and do not interfere each other; Cas9 needs a NGG PAM site and Cas12a requires a TTTN sequence for targeting^35^. Therefore, the use of Cas12a for vector control will also facilitate targeting genomic regions that we could not attack before due to restrictions with the Cas9 PAM site.

Overall, our work will provide relevant information to rapidly deploy Cas12-based strategies in disease-relevant insects. Furthermore, future applications of Cas12a-based gene drives should be applicable to different organisms, as evidenced by the plethora of CRISPR gene drives that have been successfully applied to other systems including mice, bacteria, and viruses^36–39^. Altogether, our proof-of-concept studies provide a platform that will promote further development of Cas12a-based gene-drive systems for improved insect-vector population control strategies and encourage its implementation in a wide range of organisms.

## ACKNOWLEDGMENTS

We thank Kaycie Butler and Ahana Maitra for their comments and edits on the manuscript. We thank Valentino Gantz for his scientific feedback. V.L.D.A and X.F. are authors on a patent filed by the University of California, San Diego (PCT/US2022/044487) related to the Cas12a gene-drive system described in this work.

## AUTHOR CONTRIBUTIONS

V.L.D.A and X.F conceived the project. V.L.D.A and X.F designed and obtained the plasmids and *Drosophila* transgenic lines. V.L.D.A, X.F, S.S.J, and E.O performed the experiments. V.L.D.A and S.S.J performed the figure visualizations. V.L.D.A wrote the manuscript, which was edited by all the authors.

## METHODS

### Plasmid construction

All plasmids were built following standard molecular biology approaches. Plasmids were constructed by Gibson Assembly using NEBuilder HiFi DNA Assembly Master Mix (New England BioLabs Cat. #E2621) and transformed into NEB 10-beta electrocompetent *E.coli* (New England BioLabs Cat. #3020). Plasmids were purified using a Qiagen Plasmid Midi Kit (Qiagen Cat. #12143), and DNA sequences were confirmed by Sanger sequencing.

### Generation of transgenic lines

Plasmids of Cas nucleases and the gene drive (CopyCat) were sent to Rainbow Transgenic Flies, Inc. for injection with a Cas9/Cas12a-expressing plasmid. For Cas insertions, we injected a plasmid containing the Cas nucleases and homology arms for integration along with a second plasmid expressing a gRNA targeting *yellow* for Cas12a and inserted it into the *ice2* gene in linkage with the ebony locus for Cas9 integration. CopyCat gene-drive constructs were injected into transgenic flies already expressing Cas9 or Cas12a to facilitate CopyCat insertion. The injected G_0_ larvae were sent back to us and were allowed to develop. The adult flies hatching from injected embryos were crossed to each other in pools of 3–5 males with 3–5 females. The resulting G_1_ individuals were then screened for green fluorescence in their abdomen as a marker of the CopyCat transgene insertion (*e1*- and *e4*-GDs). Cas12a insertion was identified by scoring DsRed fluorescence in the eye. Cas9 transformants were identified by scoring DsRed in the thorax. Homozygous lines for each strain were constructed from single transformants crossed to flies of the opposite sex and following the recessive body color phenotype and/or fluorescent marker in subsequent generations. Proper transgene integration in each strain was molecularly confirmed by PCR and Sanger sequencing.

### Fly rearing and crosses

Flies were fed on standard cornmeal. Fly stocks were kept at 18°C, and all experimental crosses and cages were maintained at 18°C, 25°C, or 29°C, depending on the experiment. Flies were anesthetized with CO2 when scoring phenotypes and preparing crosses. For all experimental crosses, virgin females were collected as pupae and crossed on the same day they were hatched. F0 crosses were made in pools of 3–5 males crossed to 3–5 females. F1 crosses were made in single pairs and left for five days before removing the flies (**Figs. 1,2**). After all the flies emerged, F2 flies were scored for sex, body/eye color, and fluorescence (DsRed and GFP) as a marker of transgene inheritance using a Leica M165 F2 stereomicroscope with fluorescence. All gene drive experiments were performed in a high-security ACL2 (Arthropod Containment Level 2) facility. Crosses were carried out in shatter-proof polypropylene vials (Genesee Scientific Cat. #32-120). All flies and vials were frozen for 48 hours before being removed from the facility, autoclaved, and discarded as biohazardous waste.

### Graph generation and statistical analysis

All graphs were generated using GraphPad Prism 9 and Adobe Illustrator. Statistical analyses were performed using GraphPad Prism 9. To analyze differences between *e1*-GD and *e4*-GD tested at different temperatures, two-way ANOVA and Tukey’s multiple comparisons test were performed (**Fig. 1**).

## DATA AVAILABILITY STATEMENT

The plasmid sequences for the constructs generated in this work will be shared in the GenBank database before publication. In addition, all data covering the raw phenotypical scoring data collected in the gene-drive experiments is provided in Microsoft Excel format (.xlsx), as indicated in the manuscript. All other data and information are available upon request.

