## Supplementary Figures for "Next-generation CRISPR gene-drive systems using Cas12a nuclease"

**Supplementary Figure 1**

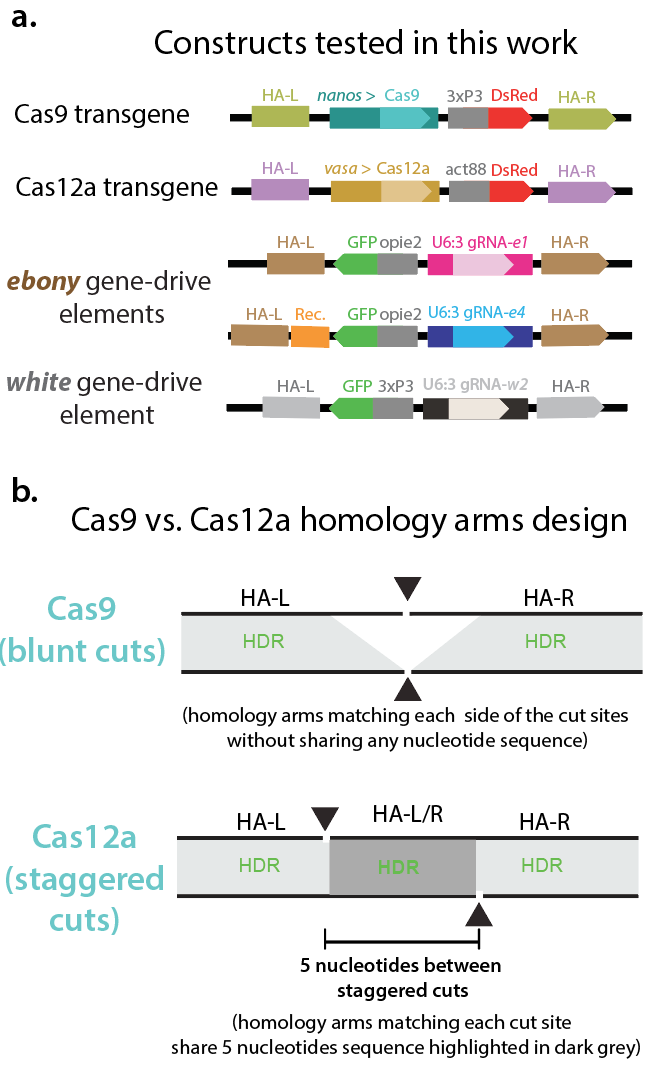

**Fig S1 - Related to Figs.1-2 –. Gene drive experiments and design. (a)** Constructs tested in this work. All transgenes were inserted by HDR and contain two homology arms (HA-L and HA-R). Cas9 is driven by the germline *nanos* promoter and marked with DsRed in the eye using the 3xP3 promoter; transgene inserted into the *ice2* locus. Cas12 is driven by the *vasa* promoter and marked with DsRed in the thorax using the opie2 promoter; transgene inserted into the *yellow* locus. Gene drive elements integrated into the *ebony* gene (e1 and e4), gRNAs driven by the *Drosophila* U6:3 promoter and marked with GFP in the abdomen. The *e4*-GD contains a DNA rescue sequence or recoded sequence (Rec.) that restores ebony function once the transgene is integrated. The *w2-*GD targeting the white gene is tagged with GFP in the eye. **(b)** Homology arms (HA) design differences between Cas9-based and Cas12a gene drives. Cas9 nuclease introduces two blunt cuts. HAs (in light grey) are designed to match each side of the cut site without any overlapping. HA-left (HA-L) and HA-right (HA-R) displayed in light green do not share any nucleotide sequence. Our Cas12a-based gene-drives inserted into *ebony* and tested in this work contain both HAs covering the gap/space (5 nucleotides) between staggered cuts. Therefore, both HA-L and HA-R contain 5 nucleotides similarity in their edge sequences highlighted in dark grey.

**Supplementary Figure 2**

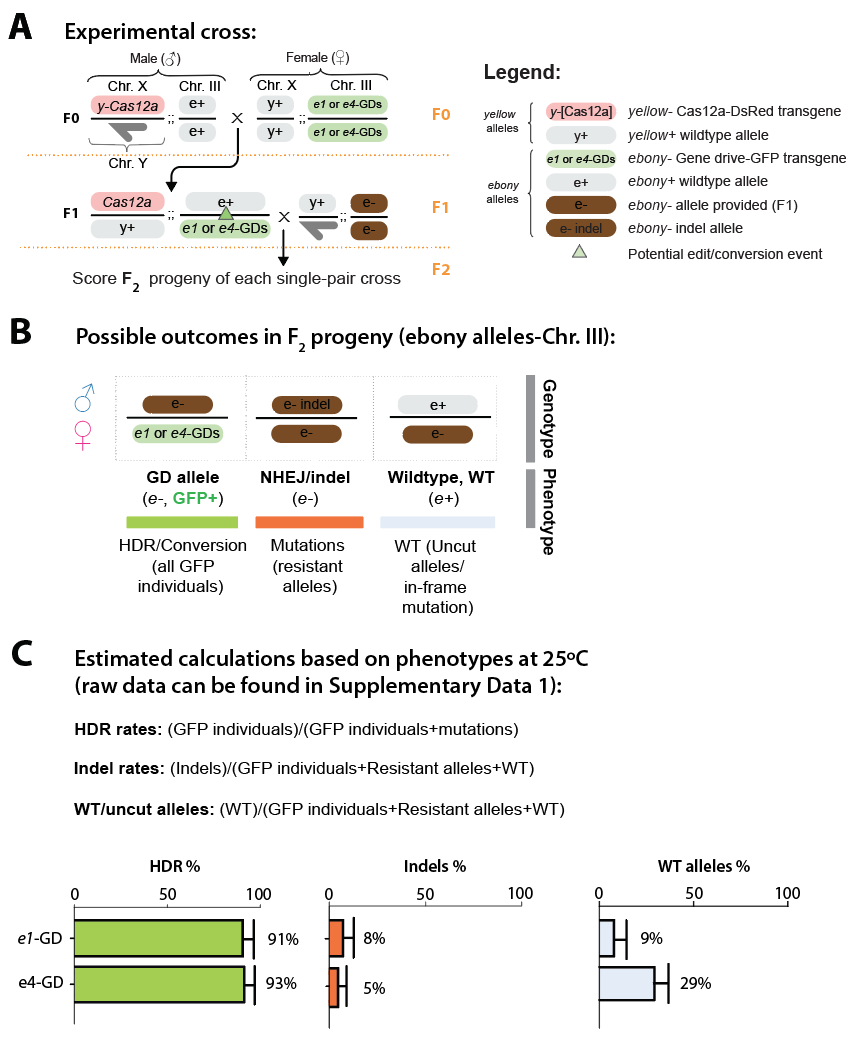

**Fig S2 - Related to Fig.1 – Estimated conversion (HDR), indels/resistant alleles and wildtype alleles produced by the Cas12-based CopyCat elements at 25°C**. **(a)** Experimental cross with alleles present in our experimental design is depicted. *Yellow* is the chromosome where the Cas12a was inserted. Our gene-drive (GD) elements were inserted in the recessive *ebony* gene. **(b)** The results were graphed according to three possible categories based on the phenotypic readouts: 1) Estimated allelic conversion or HDR (presenting as GFP) 2) indels/resistant allele events – alleles that were cut but were not converted (presenting as GFP-, and ebony phenotype), and 3) wildtype – these individuals displayed wildtype body color, and GFP-, suggesting that these alleles were not acted upon. **(c)** Graphics displaying HDR, Indels and WT alleles rates. HDR: ([all GFP individuals] / [all GFP + indels/mutations]) Indels: ([all GFP + indels] / [all GFP + indels + WT]) WT/uncut alleles: ([WT alleles] / [all GFP + indels + WT]) (see calculation/raw data on **Supplementary Data 1** for more information).
